# Composition of the Survival Motor Neuron (SMN) complex in *Drosophila melanogaster*

**DOI:** 10.1101/468223

**Authors:** A. Gregory Matera, Amanda C. Raimer, Casey A. Schmidt, Jo A. Kelly, Gaith N. Droby, David Baillat, Sara ten Have, Angus I. Lamond, Eric J. Wagner, Kelsey M. Gray

**Affiliations:** Integrative Program for Biological and Genome Sciences, The University of North Carolina, Chapel Hill, NC, 27599, USA; Curriculum in Genetics and Molecular Biology, The University of North Carolina, Chapel Hill, NC, 27599, USA; Lineberger Comprehensive Cancer Center, The University of North Carolina, Chapel Hill, NC, 27599, USA; Department of Biology and The University of North Carolina, Chapel Hill, NC, 27599, USA; Department of Genetics, The University of North Carolina, Chapel Hill, NC, 27599, USA; Department of Biochemistry and Molecular Biology, University of Texas Medical Branch, Galveston, TX 77550, USA; Centre for Gene Regulation and Expression, School of Life Sciences, University of Dundee, Dundee, DD15EH, UK

## Abstract

Spinal Muscular Atrophy (SMA) is caused by homozygous mutations in the human *survival motor neuron 1* (*SMN1*) gene. SMN protein has a well-characterized role in the biogenesis of small nuclear ribonucleoproteins (snRNPs), core components of the spliceosome. SMN is part of an oligomeric complex with core binding partners, collectively called Gemins. Biochemical and cell biological studies demonstrate that certain Gemins are required for proper snRNP assembly and transport. However, the precise functions of most Gemins are unknown. To gain a deeper understanding of the SMN complex in the context of metazoan evolution, we investigated the composition of the SMN complex in *Drosophila melanogaster.* Using a stable transgenic line that exclusively expresses Flag-tagged SMN from its native promoter, we previously found that Gemin2, Gemin3, Gemin5, and all nine classical Sm proteins, including Lsm10 and Lsm11, co-purify with SMN. Here, we show that CG2941 is also highly enriched in the pulldown. Reciprocal co-immunoprecipitation reveals that epitope-tagged CG2941 interacts with endogenous SMN in Schneider2 cells. Bioinformatic comparisons show that CG2941 shares sequence and structural similarity with metazoan Gemin4. Additional analysis shows that three other genes (CG14164, CG31950 and CG2371) are not orthologous to Gemins 6-7-8, respectively, as previously suggested. In *D.melanogaster,* CG2941 is located within an evolutionarily recent genomic triplication with two other nearly identical paralogous genes (CG32783 and CG32786). RNAi-mediated knockdown of CG2941 and its two close paralogs reveals that Gemin4 is essential for organismal viability.

## Introduction

Spinal muscular atrophy (SMA) is a pediatric neuromuscular disorder caused by mutation or loss of the human *survival motor neuron 1* (*SMN1*) gene (Lefebvre *et al.* 1995). Approximately 95% of SMA patients have homozygous deletions in *SMN1*, and the remaining ∼5% are hemizygous for the deletion over a missense mutation in *SMN1* (Burghes and Beattie 2009). Despite the great progress that has been made therapeutically, the etiology of SMA remains poorly understood. SMN’s best understood function is in the biogenesis of spliceosomal uridine-rich small nuclear ribonucleoproteins (UsnRNPs), a fundamental process important for all eukaryotic cells (Battle *et al.* 2006a; Matera *et al.* 2007; Coady and Lorson 2011; Fischer *et al.* 2011; Matera and Wang 2014). Additional tissue-specific functions for SMN have also been reported (for reviews, see Fallini *et al.* 2012; Hamilton and Gillingwater 2013; Shababi *et al.* 2014; Nash *et al.* 2016; Chaytow *et al.* 2018).

To date, no definitive link has been established between a specific function of SMN and SMA pathogenesis. Moving forward, the key question in the field is to identify which of the many potential activities of SMN lie at the root of the neuromuscular dysfunction. Given that SMA is a spectrum disorder with a broad range of phenotypes (Tizzano and Finkel 2017; Groen *et al.* 2018), it seems likely that the most severe forms of the disease would involve loss of more than one SMN-dependent pathway. Thus, understanding the molecular etiology of the disease is not only important for the basic biology, but also for targeting and refining therapeutic strategies (Groen *et al.* 2018; Sumner and Crawford 2018).

As outlined in Fig. 1, SMN works in close partnership with a number of proteins, collectively called Gemins (Paushkin *et al.* 2002; Shpargel and Matera 2005; Otter *et al.* 2007; Borg and Cauchi 2013). The N-terminal domain of SMN interacts with a protein called Gemin2 (Liu *et al.* 1997; Wang and Dreyfuss 2001), whereas the C-terminal region contains a YG-zipper motif (Martin *et al.* 2012) that drives SMN self-oligomerization, as well as binding to Gemin3 (Lorson *et al.* 1998; Pellizzoni *et al.* 1999; Praveen *et al.* 2014; Gupta *et al.* 2015). Gemin2 (Gem2) heterodimerizes with SMN and is the only member of the complex that is conserved from budding yeast to humans (Fischer *et al.* 1997; Kroiss *et al.* 2008). When provided with *in vitro* transcribed Sm-class snRNAs, purified recombinant SMN and Gem2 are sufficient for Sm core assembly activity (Kroiss *et al.* 2008). *In vivo*, Gemin5 (Gem5) is thought to play an important role in snRNA substrate recognition (Battle *et al.* 2006b; Bradrick and Gromeier 2009; Lau *et al.* 2009; Yong *et al.* 2010; Jin *et al.* 2016), but it has also been independently identified as a cellular signaling factor (Gates *et al.* 2004; Kim *et al.* 2007) and a translation factor (reviewed in Pineiro *et al.* 2015).

**Figure 1.**
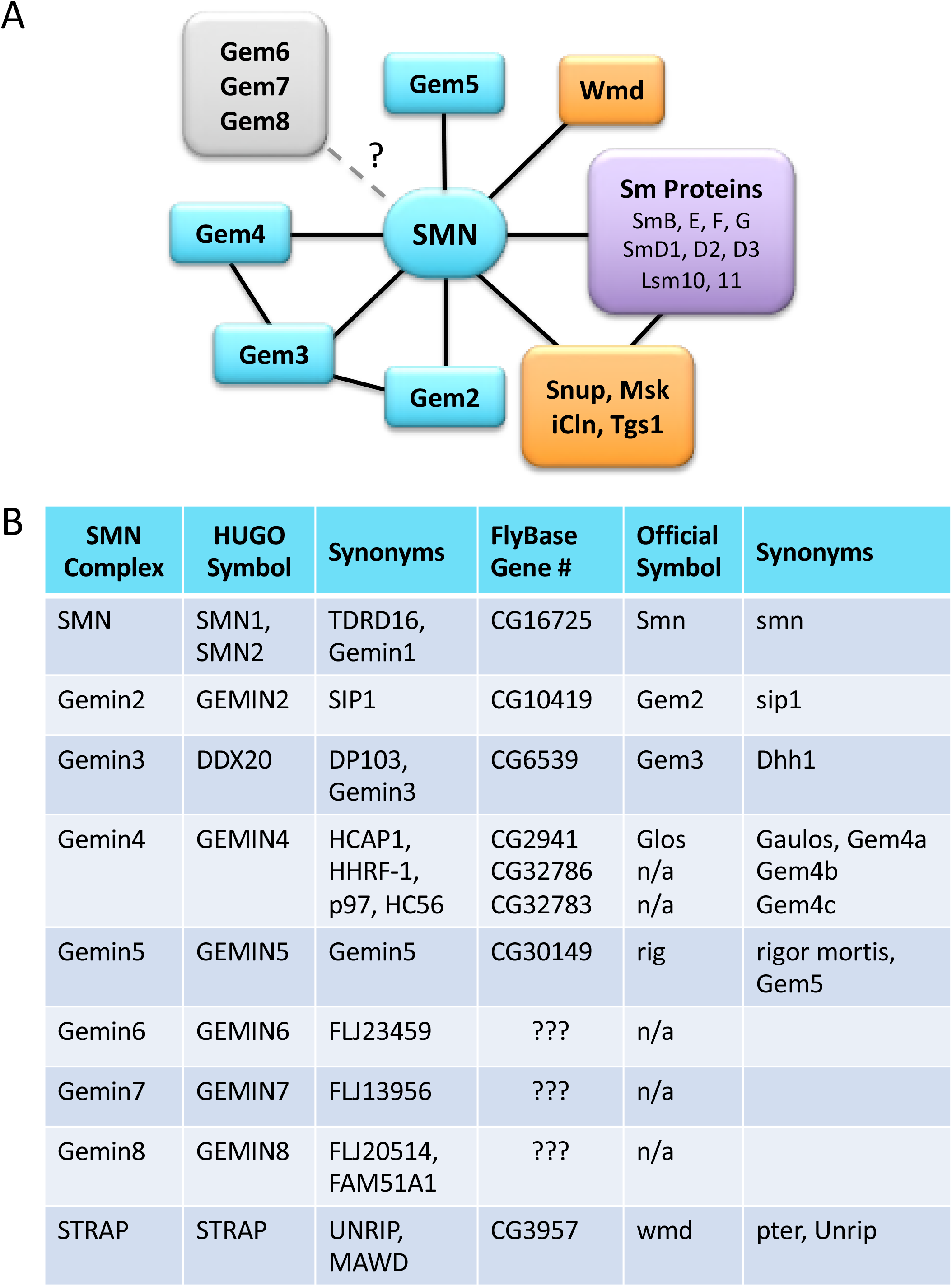
Protein interaction network of the metazoan SMN complex. (**A**) In addition to the core Gemin family members (shown in teal), SMN is known to form complexes with various Sm protein substrates (purple) and RNP biogenesis factors (orange). Snup, Msk, iCln, Tgs1 and Wmd, correspond to SPN1/Snurportin1, Moleskin/Importin7, CLNS1A/Chloride channel nucleotide-sensitive 1A, TGS1/Trimethylguanosine synthase 1, and STRAP/Unrip, respectively. The presence of three of the Gemins (Gem6, Gem7 and Gem8) within dipteran genomes is uncertain (?). (**B**) Reference table comparing the names, symbols and synonyms of SMN complex members in humans and flies.

Gemin3/Dp103 (Gem3) is a DEAD-box helicase (Charroux *et al.* 1999; Grundhoff *et al.* 1999) that interacts with the SMN•Gem2 hetero-oligomer and is reported to play roles in transcriptional repression (Yan *et al.* 2003) and microRNA activity (Mourelatos *et al.* 2002) in addition to its role in Sm-core assembly (Shpargel and Matera 2005), reviewed in (Curmi and Cauchi 2018). Gemin4 (Gem4) is tethered to the SMN complex via direct binding to Gem3 (Charroux *et al.* 2000; Meister *et al.* 2000). Gem4 has been implicated in nuclear receptor binding (Di *et al.* 2003; Yang *et al.* 2015), microRNA biology (HutvÁgner and Zamore 2002; Meister *et al.* 2005) and in nuclear import of the SMN complex (Narayanan *et al.* 2004; Meier *et al.* 2018). In human cell lysates, SMN and Gemins2-4 are thought to be essential for proper assembly of the Sm core (Shpargel and Matera 2005). Consistent with this notion, complete loss-of-function mutations in *Smn, Gem2, Gem3* and *Gem4* in mice result in embryonic lethality (Schrank *et al.* 1997; Jablonka *et al.* 2002; Mouillet *et al.* 2008; Meier *et al.* 2018). Partial loss-of-function mutations in genes encoding the Gemins have not been reported. Functions of the other members of the SMN complex are largely unknown.

Gemins6-8 (Gem6-7-8) and STRAP (serine-threonine kinase receptor associated protein, a.k.a. UNRIP, UNR-interacting protein; or Wmd/CG3957, wing morphogenesis defect) are peripheral members of the SMN complex and may thus serve in a regulatory capacity. In support of this idea, STRAP/UNRIP is found in a separate complex with a cap-independent translation factor called UNR (Hunt *et al.* 1999) and has been shown to modulate its function (Carissimi *et al.* 2005; Grimmler *et al.* 2005). STRAP binds to the SMN complex via an interaction with Gem7 (Otter *et al.* 2007). In *Drosophila*, mutations in the STRAP orthologue, Wmd/CG3957, cause defects in development of the adult wing (Khokhar *et al.* 2008), suggesting that this protein may not be essential for basal assembly of spliceosomal snRNPs. Gem6-7-8 forms a subcomplex that is tethered to SMN via interaction with Gem8 (Carissimi *et al.* 2006; Otter *et al.* 2007). Gem6 and Gem7 are Sm-like proteins that heterodimerize with one another, but the roles played by these factors in snRNP biogenesis are unknown. Mutations in Gem6-7-8 have yet to be described.

In *Drosophila*, the SMN complex (as defined by proteins that stably co-purify with SMN) was originally thought to contain only Gem2 and Gem3 (Kroiss *et al.* 2008). Bioinformatic analysis suggested that the gene *rigor mortis* (Rig) encodes a potential metazoan *Gemin5* orthologue, but Gem5/Rig protein failed to co-purify with SMN in Schneider2 (S2) cells (Kroiss *et al.* 2008). Notably, transgenic expression of tagged Gem2 and Gem5/Rig constructs showed that these two proteins colocalize with endogenous SMN in cytoplasmic structures called U bodies (Cauchi *et al.* 2010). Consistent with the cytological studies, we found that Gem5/Rig co-purified with SMN in *Drosophila* embryos that were engineered to exclusively express Flag-tagged SMN (Gray *et al.* 2018). However, *Drosophila* Gem6-7-8 proteins were neither identified bioinformatically nor were they shown to biochemically coprecipitate with SMN in S2 cells (Kroiss *et al.* 2008).

A recent study presented evidence suggesting the “full conservation” of the SMN complex in fruit flies (Lanfranco *et al.* 2017), and arguing that Gem4 and Gem6-7-8 are present in Diptera. In this report, we re-investigate this issue, showing that among the four novel factors identified by Lanfranco et al. (2017), only Gaulos (CG2941) is orthologous to a metazoan SMN complex protein (Gem4). Using comparative genomic analysis, we conclusively demonstrate that Hezron (CG14164), Sabbat (CG31950) and Valette (CG2371) are not orthologous to metazoan Gemin6, Gemin7 and Gemin8, respectively. The genes encoding these three *Drosophila* proteins are actually orthologous to three distinct and highly conserved metazoan genes. The implications of these findings are discussed. Furthermore, we purified SMN complexes from *Drosophila* embryos and found that endogenous Gaulos co-precipitates with SMN but Valette, Sabbat and Hezron do not. Phylogenetic analysis demonstrates that *Gaulos/CG2941* is actually part of a genomic triplication involving two other nearly identical gene copies, *CG32783* and *CG32786*. We interrogate the function of these three factors *in vivo* using RNA interference analysis, revealing that the function of these redundant genes is essential in flies.

## Materials and Methods

### Fly stocks

RNAi lines were obtained from the Bloomington TRIP collection. The identifying numbers listed on the chart are stock numbers. Each of the RNAi constructs is expressed from one of five VALIUM vectors and requires Gal4 for expression. All stocks were cultured on molasses and agar at room temperature (25°C).

### Antibodies and Western blotting

Embryonic lysates were prepared by crushing the animals in lysis buffer (50mM Tris-HCl, pH 7.5, 150 mM NaCl, 1mM EDTA, 1% NP-40) with 1X protease inhibitor cocktail (Invitrogen) and clearing the lysate by centrifugation at 13,000 RPM for 10 min at 4ºC. S2 cell lysates were prepared by suspending cells in lysis buffer (50mM Tris-HCl, pH 7.5, 150 mM NaCl, 1mM EDTA, 1% NP-40) with 10% glycerol and 1x protease inhibitor cocktail (Invitrogen) and disrupting cell membranes by pulling the suspension through a 25 gauge needle (Becton Dickinson). The lysate was then cleared by centrifugation at 13,000 RPM for 10 min at 4ºC. Cell fractionation was performed using a standard protocol (West *et al.* 2008). In brief, following centrifugation, cytoplasmic extracts were taken from the top 0.2mL and the nuclear pellet was resuspended in 0.2mL RIPA buffer. Western blotting on lysates was performed using standard protocols. Rabbit anti-dSMN serum was generated by injecting rabbits with purified, full-length dSMN protein (Pacific Immunology Corp, CA), and was subsequently affinity purified. For Western blotting, dilutions of 1 in 2,500 for the affinity purified anti-dSMN, 1 in 10,000 for monoclonal anti-FLAG (Sigma) were used.

### Immunoprecipitation

Lysates were incubated with Anti-FLAG antibody crosslinked to agarose beads (EZview Red Anti-FLAG M2 affinity gel, Sigma) for 2h-ON at 4C with rotation. The beads were washed with RIPA lysis buffer or three times and boiled in SDS gel-loading buffer. Eluted proteins were run on an SDS-PAGE for western blotting.

### Drosophila embryo protein lysate and mass spectrometry

0-12h Drosophila embryos were collected from Oregon-R control and Flag-SMN flies, dechorionated, flash frozen, and stored at −80°C. Embryos (approx. 1gr) were then homogenized on ice with a Potter tissue grinder in 5 mL of lysis buffer containing 100mM potassium acetate, 30mM HEPES-KOH at pH 7.4, 2mM magnesium acetate, 5mM dithiothreitol (DTT) and protease inhibitor cocktail. Lysates were centrifuged twice at 20,000 rpm for 20min at 4°C and dialyzed for 5h at 4°C in Buffer D (HEPES 20mM pH 7.9, 100mM KCl, 2.5 mM MgCl_2_, 20% glycerol, 0.5 mM DTT, PMSF 0.2 mM). Lysates were clarified again by centrifugation at 20000 rpm for 20 min at 4C. Lysates were flash frozen using liquid nitrogen and stored at −80C before use. Lysates were then thawed on ice, centrifuged at 20000 rpm for 20 min at 4C and incubated with rotation with 100 uL of EZview Red Anti-FLAG M2 affinity gel (Sigma) for 2h at 4C. Beads were washed a total of six times using buffer with KCl concentrations ranging from 100mM to 250mM with rotation for 1 min at 4°C in between each wash. Finally, Flag proteins were eluted 3 consecutive times with one bed volume of elution buffer (Tris 20mM pH 8, 100 mM KCl, 10% glycerol, 0.5 mM DTT, PMSF 0.2 mM) containing 250ug/mL 3XFLAG peptide (sigma). The eluates were used for mass spectrometric analysis on an Orbitrap Velos instrument, fitted with a Thermo Easy-spray 50cm column.

### Viability and Larval Locomotion

Males containing RNAi constructs were crossed to virgin females containing one of the Gal4 constructs balanced by CAG (Tub-Gal4) or TM6BGFP (Da-Gal4). Embryos were collected on molasses agar plates and sorted into vials using lack of GFP fluorescence. A maximum of 50 larvae were sorted into each vial. Viability was assessed based on the number of pupated or eclosed individuals compared to the starting number of larvae in each vial.

To assess larval locomotion, five wandering third instar larvae were set on a large molasses agar plate and placed in a recording chamber. Their crawling movements were recorded for at least 1 min on a digital camera (smartphone) at minimum zoom. Four recording were take for each set of larvae; at least 30 larvae total were recorded for each cross. The videos were transferred to a PC and converted to AVI files using ffmpeg (https://www.ffmpeg.org/) . The videos were then opened and converted to binary frames using Fiji/ImageJ. The wrMTrck plugin (http://www.phage.dk/plugins/wrmtrck.html) for ImageJ was used to assess the average speed of each larvae normalized to their body size (body lengths/second or BLPS).

### Northern blotting and RT-PCR

Early third instar larvae (73-77 hours post egg-laying) were homogenized in TRIzol reagent (Invitrogen) and RNA was isolated according to the manufacturer’s protocol with the following modifications: a second chloroform extraction was performed, and RNA was precipitated with 0.1 volumes of sodium acetate and 2.5 volumes of ethanol rather than isopropanol. For Northern blotting, 2500 ng of total RNA was separated on Novex 10% TBE-Urea gels (Invitrogen). RNA was transferred to GeneScreen Plus Hybridization Transfer Membrane (PerkinElmer). Blots were dried, UV cross-linked, and pre-hybridized with Rapid-hyb Buffer (GE Healthcare). Probes were prepared by 5’-end labeling oligonucleotides with [γ-^32^P]ATP (PerkinElmer) using T4 PNK (NEB). The oligonucleotide probe sequences are as follows:

U1: 5′-GAATAATCGCAGAGGTCAACTCAGCCGAGGT-3′

U2: 5′-TCCGTCTGATTCCAAAAATCAGTTTAACATTTGTTGTCCTCCAAT-3′

U4: 5′-GGGGTATTGGTTAAAGTTTTCAACTAGCAATAATCGCACCTCAGTAG-3′

U5: 5′-GACTCATTAGAGTGTTCCTCTCCACGGAAATCTTTAGTAAAAGGC-3′

U6: 5′-CTTCTCTGTATCGTTCCAATTTTAGTATATGTTCTGCCGAAGCAAGA-3′

U11: 5′-TCGTGATCGGAAACGTGCCAGGACG-3′

U12: 5′-GCCTAGAAGCCAATACTGCCAAGCGATTAGCAAG-3′

U4atac: 5′-AGCAATGTCCTCACTAGACGTTCATTGAACATTTCTGCT-3′

U6atac: 5′-CCTAGCCGACCGTTTATGTGTTCCATCCTTGTCT-3′

Following the PNK reaction, probes were purified using Microspin G-50 Columns (GE Healthcare). For hybridization, the blots were probed with the labeled oligonucleotides at 65°C. The blots were then washed twice each in 2X SSC and 0.33X SSC at 60°C. Blots were exposed to storage phosphor screens (GE Healthcare) and analyzed with an Amersham Typhoon 5 (GE Healthcare).

To analyze knockdown efficiency, total RNA was treated with TURBO DNase (Invitrogen). Following a phenol/chloroform purification, 350 ng of RNA was converted to cDNA using the SuperScript III First-Strand Synthesis System (Invitrogen). Primers for PCR are as follows:

5S_F: 5′-GCCAACGACCATACCACGCTGAA-3′

5S_R: 5′-AACAACACGCGGTGTTCCCAAGC-3′

CG2941_F: 5′-TGTGGTATTGGCAGGACGGTCT-3′

CG2941_R: 5′-CCTTGTGCTTCAATTTGCTCACTTGGTT-3′

Triplicate_F: 5′-CCAGATAGCCTGCATGGAACATCG-3′

Triplicate_R: 5′-CTCCCGCTTTAATGGATCATTGAGGG-3′

### Data availability statement

All fly strains, probe sequences and plasmids are available upon request. The authors affirm that all data necessary for confirming the conclusions of the article are present within the article, figures, and tables.

## Results and Discussion

To identify *Drosophila melanogaster* proteins that might correspond to those known to be contained within the human SMN complex (Fig. 1B), we first carried out *in silico* bioinformatic analyses. As mentioned above, there have been conflicting reports regarding the conservation (or lack thereof) of certain Gemin proteins in *Drosophila* (Kroiss *et al.* 2008; Lanfranco *et al.* 2017). In particular, Gem4, Gem6, Gem7 and Gem8 were originally thought to have been lost from Dipteran genomes, although clear orthologues of Gem6-7-8 can been readily identifed in Hymenoptera, and other insects (e.g. search https://www.ncbi.nlm.nih.gov/protein).

### Hezron (CG14164) is orthologous to Lsm12, not Gemin6

Lanfranco et al. (2017) recently suggested that *Hezron/CG14164* encodes the *Drosophila* orthologue of Gem6. Our bioinformatic analysis suggested that CG14164 is actually more closely related to human Lsm12, an Sm-like protein that contains a C-terminal methyltransferase domain (Albrecht and Lengauer 2004). Because Gem6 is also an Sm-like protein (Gem6, Gem7 and Lsm12 are members of the Sm protein superfamily), we carried out a side-by-side comparison of both Lsm12 and Gem6 proteins from a variety of vertebrates and invertebrates (Fig. 2). That is, we selected Lsm12 and Gem6 protein pairs from each of five different species (human, chicken, fish, bug and wasp) and aligned them together to identify highly conserved diagnostic amino acid residues in each orthology cluster. Notably, there are two very closely related proteins in *D. melanogaster*, CG14164 and CG15735, both of which were included in this comparison. As shown in Fig. 2, CG14164/Hezron is clearly more closely related to the Lsm12 cluster than it is to that of Gem6 (compare diagnostic Lsm12 and Gem6 residues shaded in red and blue, respectively). In nearly every case, CG14164 tracks with the Lsm12 sequences, including the locations of conserved insertions and deletions. Therefore, we conclude that *Hezron/CG14164* is orthologous to metazoan *Lsm12* and that the ancestral *Gem6* gene has been lost in *Drosophila*.

**Figure 2.**
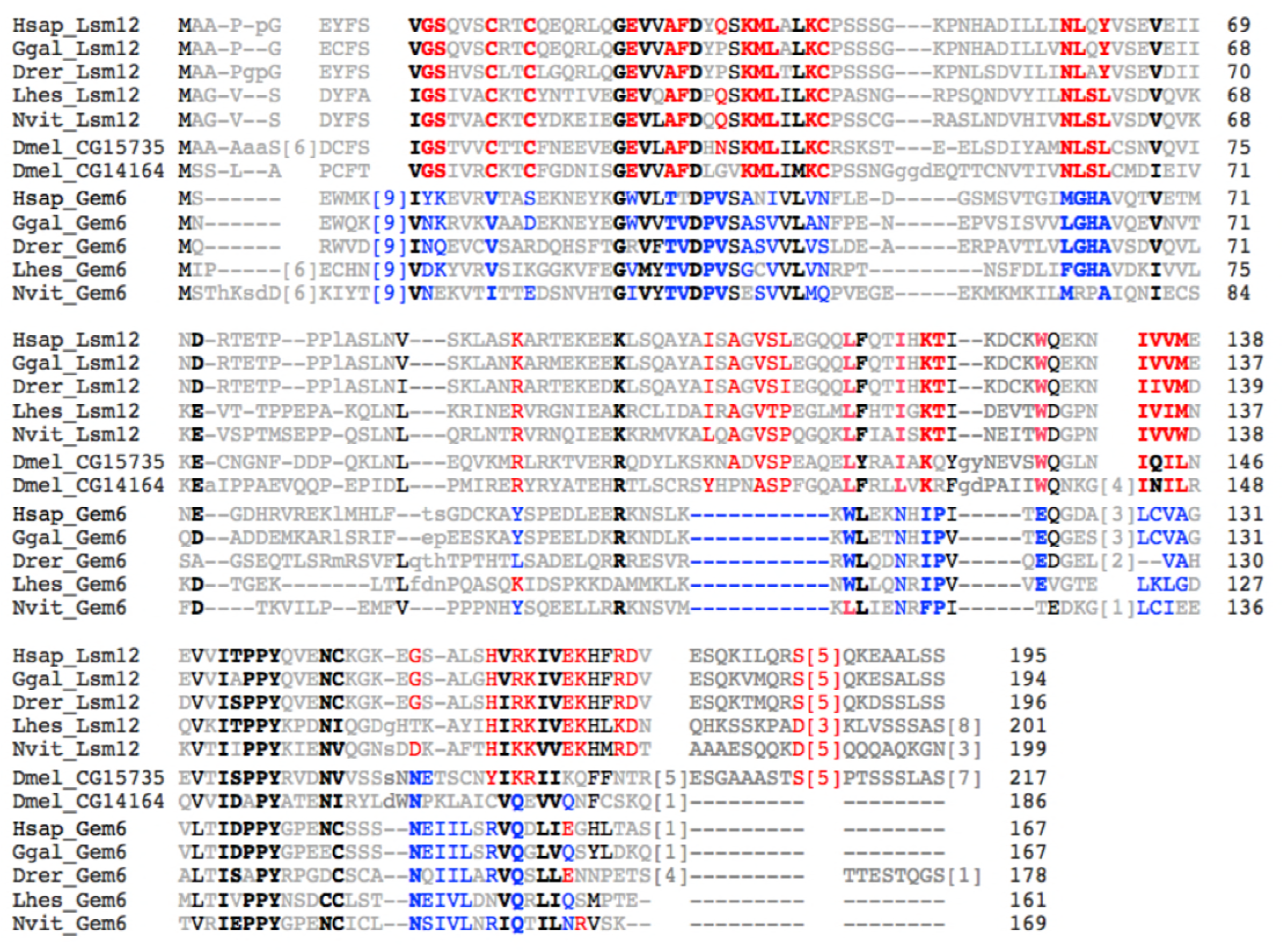
CG14164/Hezron is orthologous to Lsm12, not Gemin6. Amino acid alignment of Lsm12 and Gemin6 (Gem6) protein sequences from a variety of metazoan species, including: *Homo sapiens* (Hsap), *Gallus gallus* (Ggal), *Danio rerio* (Drer), *Lygus hesperus* (Lhes), and *Nasonia vitripennis* (Nvit). Two paralogous *D. melanogaster* proteins CG15735 and CG14164, are shown for comparison. Each of the fruitfly proteins shows a high degree of similarity to Lsm12, as compared to Gem6. Residues shaded in red are conserved in Lsm12 orthologs and those shaded in blue are conserved in the Gem6 orthologs.

CG15735 was recently shown to function as an Ataxin-2 adaptor (Lee *et al.* 2017); a genetic analysis of CG14164 has not been reported. We note that CG15735/Lsm12a (217aa) is slightly longer than Hezron/CG14164/Lsm12b (186aa) and that the two proteins begin to diverge only at their respective C-termini (Fig. 2). There is the barest hint of similarity between CG14164 and Gem6 in this region. It is tempting to speculate that an ancestral recombination between *Gem6* and *Lsm12* might have created *Hezron/CG14164*. Additional experiments will be required in order to address these evolutionary relationships, as well as to determine whether or not Hezron/CG14164/Lsm12b protein might have been co-opted into the extant *Drosophila* SMN complex (see below).

### Sabbat (CG31950) is orthologous to Naa38, not Gemin7

CG31950/Sabbat was also identified by Lanfranco et al. (2017) as a potential orthologue to Gem7 and member of the *Drosophila* SMN complex. We found that CG31950 was, in fact, more similar to an N-terminal acetyltransferase auxiliary subunit, Naa38 (Varland *et al.* 2015; Aksnes *et al.* 2016). An alignment of Naa38 proteins from human, fish, honeybee, sea urchin and fission yeast, along with the Gem7 orthologues from these same five species reveals that CG31950 is much more similar to the Naa38 orthology cluster than it is to that of Gem7 (Fig. 3). For purposes of comparison, we also include in the alignment the most closely related protein in a second fruitfly genome, *D. hydei* (XP_023177506.1), along with CG31950 from *D. melanogaster* (Fig. 3). Although the two clusters of proteins share an overall sequence similarity (indeed, human Naa38 is also known to contain an Sm-like fold; see https://www.uniprot.org/uniprot/I3L310) diagnostic residues within the Naa38 orthologues are shared by CG31950. In contrast, the highly conserved regions of Gem7, including the relative positions of insertions and deletions, do not track with CG31950 (see shaded residues in Fig. 3). Hence, we conclude that Sabbat/CG31950 is orthologous to Naa38.

**Figure 3.**
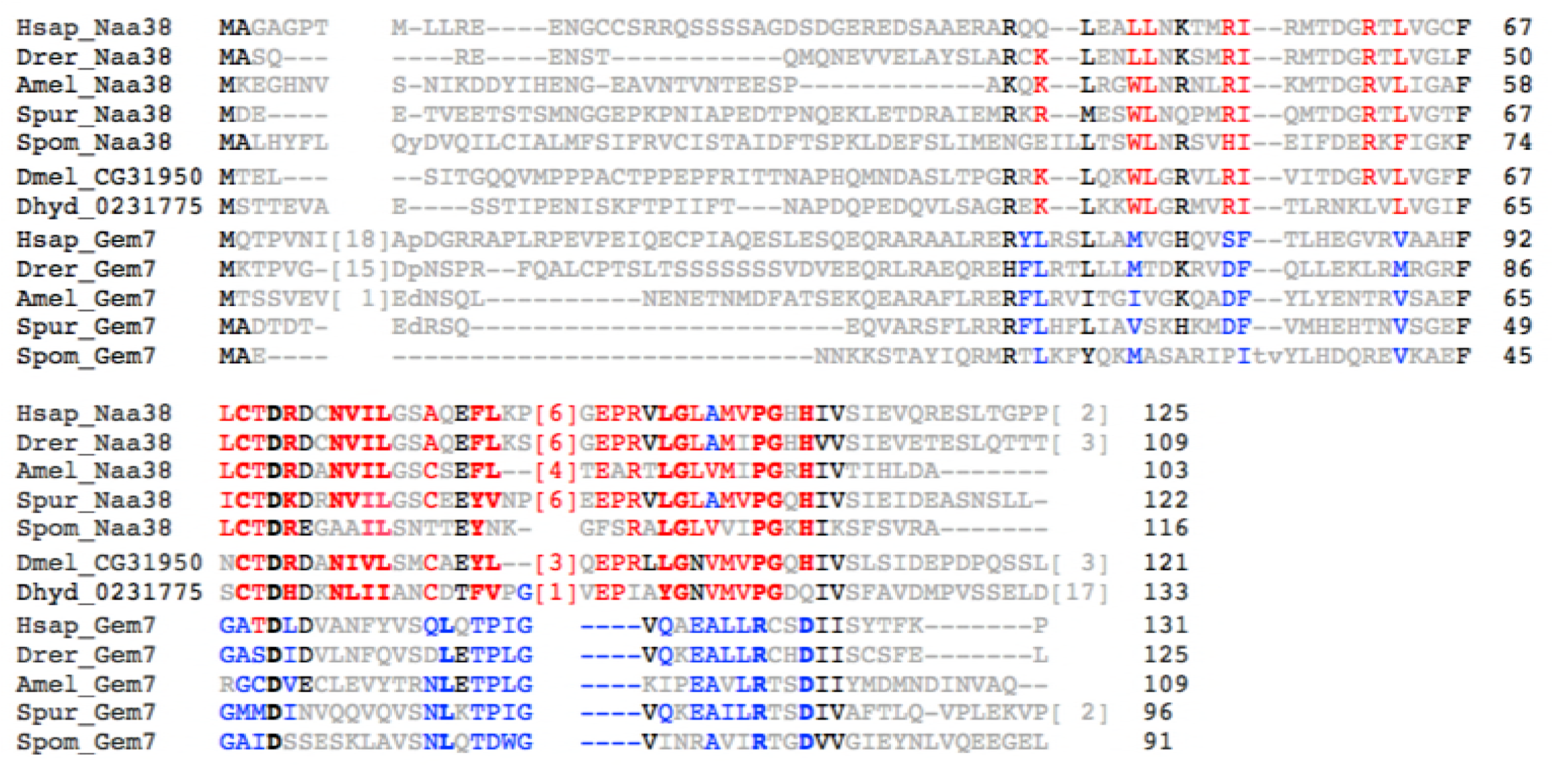
CG31950/Sabbat is orthologous to Naa38, not Gemin7. Amino acid alignment of Gemin7 (Gem7) and Naa38 protein sequences from a variety of metazoan species, including: *Homo sapiens* (Hsap), *Danio rerio* (Drer), *Apis mellifera* (Amel), *Strongylocentrotus purpuratus* (Spur) and *Schizosaccharomyces pombe* (Spom). Two orthologous fruitfly proteins from *D. melanogaster* (Dmel_CG31950) and *D. hydei* (Dhyd_0231775) are shown for comparison. Residues in red are conserved in Naa38 sequences and those in blue are conserved in Gem7.

### Valette (CG2371) is orthologous to CommD10, not Gemin8

Similar to the situation with Gem6 and Gem7, we found that CG2371, identified by Lanfranco et al. (2017) as the potential Gem8 ortholog, is more closely related to a protein called CommD10 (Fig. 4). In humans, there are ten Comm domain paralogs, five of which are conserved in insects (Maine and Burstein 2007). These proteins are characterized by a conserved C-terminal ∼80 aa region called the Comm domain (see discussion below). Gemin8 orthologs are also conserved at their C-termini, but the structure and function of this protein is largely unknown. As shown in Fig. 4, *D.melanogaster* CG2371 and *D.yakuba* GE17608 proteins are most similar to CommD10 orthologues as compared to the Gem8 orthologs from a variety of metazoan species. The conservation of diagnostic amino acid residues between CG2371 and CommD10 (Fig. 4, shaded residues) leaves little doubt as to the ancestral relationship. Again, the interesting question of whether Valette/CG2371 might have compensated for loss of Gem8 within the *Drosophila* lineage is discussed below.

**Figure 4.**
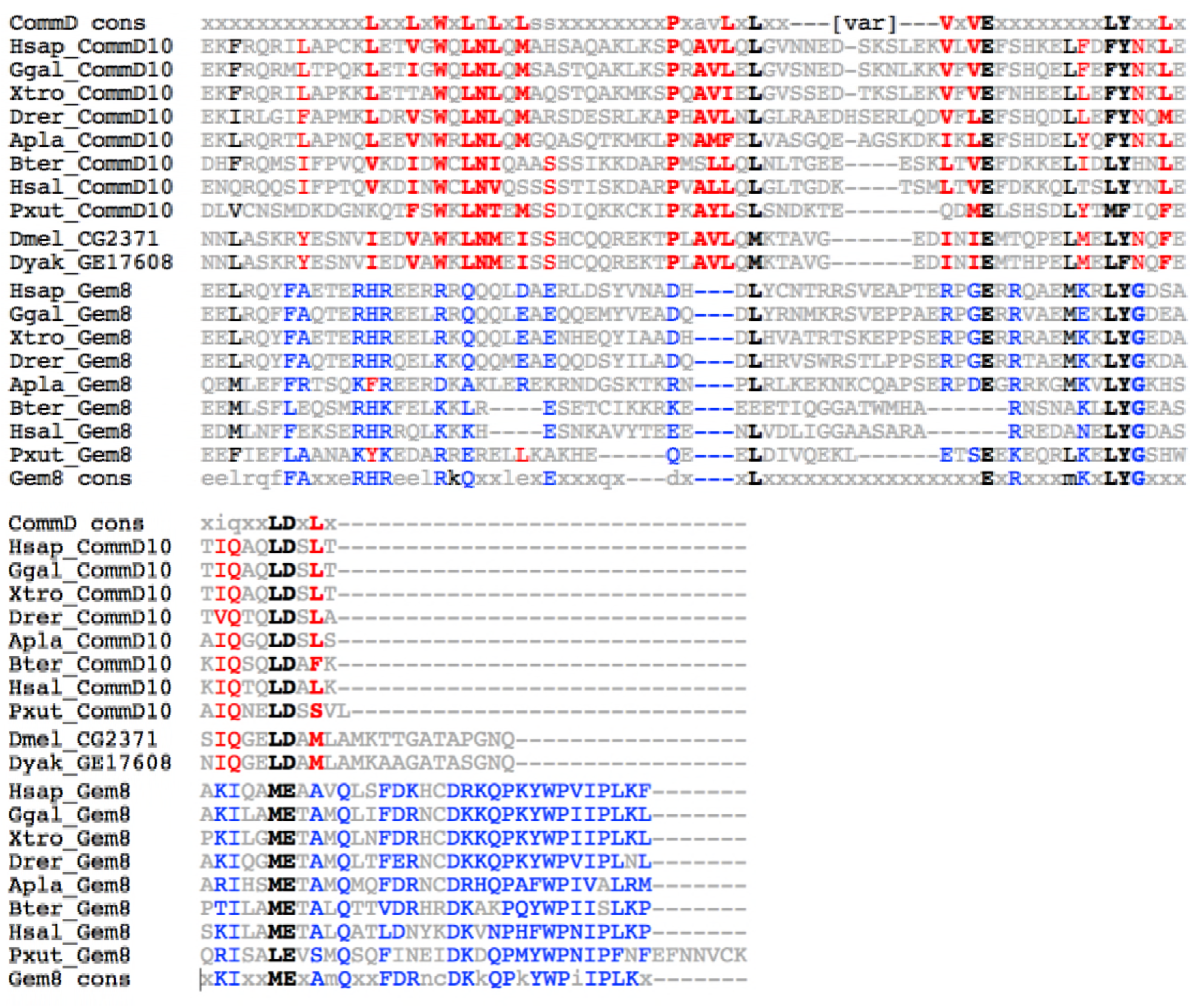
CG2371/Valette is orthologous to CommD10, not Gemin8. Amino acid alignment of the C-terminal domains of CommD10 and Gemin8 (Gem8) from a variety of metazoan species, including: *Homo sapiens* (Hsap), *Gallus gallus* (Ggal), *Xenopus tropicalis* (Xtro), *Danio rerio* (Drer), *Acanthaster planci* (Apla), and *Bombus terrestris* (Bter), *H saltator* (Hsal), and *Papilio xuthus* (Xut). Two orthologous fruitfly proteins from *D. melanogaster* (Dmel_CG2371) and *D. yakuba* (Dyak_GE17608) are shown for comparison. Red residues are conserved in CommD10 sequences whereas blue residues are conserved in Gem8. For additional comparison, consensus sequences for the C-terminal Comm domain (CommD cons) and C-terminus of Gemin8 (Gem8 cons) are shown along the top and bottom of the alignment, respectively.

### Gaulos (CG2941) is orthologous to metazoan Gemin4

In contrast to Gem6-7-8, Gem4 was originally thought to be lost from insects entirely (Kroiss *et al.* 2008), however Lanfranco et al. (2017) identified CG2941/Gaulos as a potential Gem4 orthologue. To investigate this issue, we carried out PSI-BLAST (Position-Specific Iterative Basic Local Alignment Search Tool) analysis (Altschul *et al.* 1997). Using vertebrate Gem4 proteins as seed sequences, this procedure readily identified several candidates among the Hymenoptera, but not within the Diptera (or it may take more than six iterations to converge). Anecdotally, we have found that the entire SMN complex is well preserved in many Hymenopteran genomes, and so we used the putative Gem4 sequence XP_003401506.1 from *Bombus terrestris* (Buff-tailed bumble bee) as the starting point for PSI-BLAST. We found that this seed sequence readily identified both vertebrate and invertebrate orthologs, along with three nearly identical *D. melanogaster* proteins, including CG2941, CG32783 and CG32786. An alignment of a subset of these identified proteins is presented in Fig. S1. As shown, the overall conservation of Gem4 is rather modest. For comparison, the putative Gem4 orthologue (GH21356) from a distantly related fruitfly, *D. grimshawi*, is also shown. Notably, an analysis of the predicted secondary structure of Gem4 orthologs shows a high degree of conservation (not shown). Thus, despite the fact that the three *D. melanogaster* proteins (CG2941, CG32786 and CG32783) are most closely related to metazoan Gem4, it is difficult to assign orthology on the basis of amino acid conservation alone.

Importantly, Lanfranco et al. (2017) did not base their conclusions solely on bioinformatics; these authors also showed that CG2941 interacts with Gem3 in a targeted genetic modifier screen. Flies expressing low-levels of a dominant-negative *Gem3* construct lacking its N-terminal helicase domain, called *Gem3^BART^* (Borg *et al.* 2015), were used in combination with a deficiency allele *Df(1)ED6716* (Ryder *et al.* 2007) that spans the 3F2-4B3 interval on the X chromosome that includes *CG2941/Gaulos* (Lanfranco *et al.* 2017). Loss of one copy of this region in combination with pan-muscular expression of *Gem3^BART^* led to a marked age-dependent enhancement of the phenotype (Lanfranco *et al.* 2017). These results were encouraging because Gem4 was originally identified as a putative Gem3 cofactor (Charroux *et al.* 2000), and so a reduction in *Gem4* gene copy-number might reasonably be expected to enhance the phenotype of a *Gem3* hypomorph.

Furthermore, a systematic coaffinity purification analysis of the *Drosophila* proteome showed that CG2941 is capable of forming a complex in S2 cells transfected with HA-tagged SMN (Guruharsha *et al.* 2011). Similarly, co-transfection of S2 cells with HA-tagged SMN and GFP-tagged CG2941/Gaulos cells confirmed this interpretation (Lanfranco *et al.* 2017). Previously, we carried out proteomic profiling of embryonic lysates from transgenic flies expressing Flag-SMN (Gray *et al.* 2018). Notably, these animals express SMN protein from the native *Smn* control regions in an otherwise null background (Praveen *et al.* 2012; Gray *et al.* 2018). In order to conclusively determine whether CG2941 forms a complex with SMN under endogenous conditions, we directly analyzed eluates of this purification by label-free mass spectrometry (Fig. 5A). We also carried out SDS-PAGE analysis of the purified samples, followed by silver staining (Fig. 5B). In addition to identifying all of the known Sm protein substrates of SMN, we also identified Gem2, Gem3, Gem5 and CG2941 as highly-enriched SMN binding partners (Fig. 5). Notably, mass spectrometry failed to detect CG14164 (Hezron/Lsm12b), CG31950 (Sabbat/Naa38) or CG2371 (Valette/CommD10) among the co-purified proteins (see below for a discussion).

**Figure 5.**
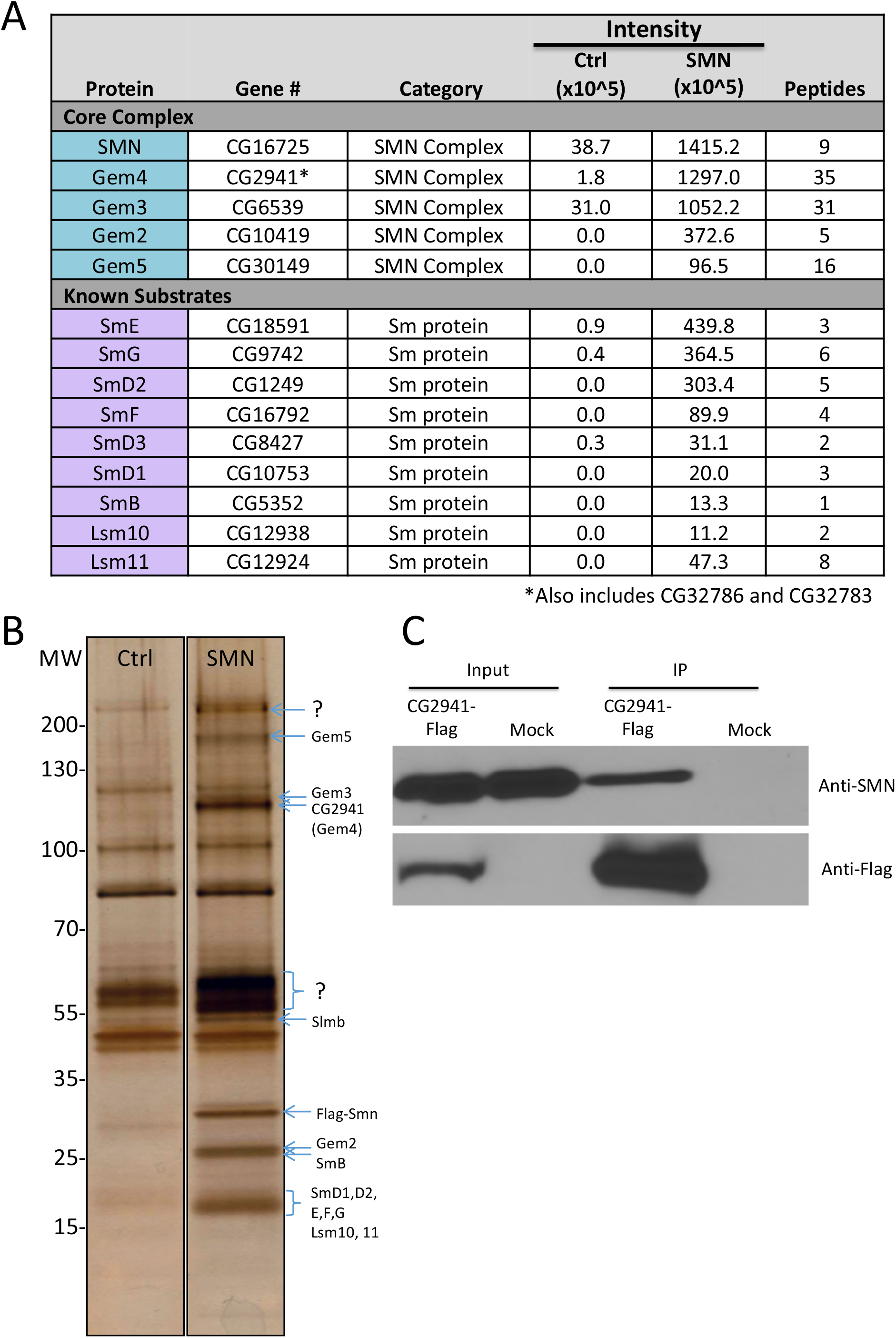
Flag-SMN immunopurified lysates contain members of the known SMN complex and known substrates. (**A**) Flag-purified eluates were analyzed by label-free mass spectrometry. Proteins in the core SMN complex and known substrates were highly enriched in the SMN sample as compared with Ctrl. (**B**) Lysates from Oregon-R control (Ctrl) *Drosophila* embryos and those that exclusively express Flag-SMN (SMN) were immunopurified. Protein eluates were separated by SDS-PAGE and silver stained. Band identities were predicted by size. (**C**) Following transfection of CG2941-Flag in *Drosophila* S2 cells and immunoprecipitation (IP) of total cell lysates with anti-Flag beads, western analysis was carried out using anti-SMN antibodies. Mock transfected cells were analyzed in parallel. Co-purification of endogenous SMN was detected in the CG2941-Flag IP lane but not in the control lane (Mock).

We also note that CG32786 and CG32783 are so similar to CG2941 that most of their tryptic peptides are indistinguishable from one another. However, we identified 35 peptides corresponding to these three proteins and their overall enrichment in the purified eluates was comparable to that of the other core members of the SMN complex (Fig. 5A). To confirm this interaction, we performed a reciprocal coimmunoprecipitation analysis with CG2941-Flag in S2 cells. As shown in Fig. 5C, CG2941-Flag also co-purifies with endogenous SMN. On the basis of these findings, we conclude that CG2941/Gaulos is indeed orthologous to human Gem4.

### CG2941 is ancestral to CG32786 and CG32783 and is part of a genomic triplication

As shown in Fig. 6A, CG2941, CG32786 and CG32783 are tightly linked in the 3F7-3F9 interval on the *D. melanogaster* X chromosome. A comparison of their DNA sequences reveals that these three genes are extremely similar, with CG32786 and CG32783 being more closely related to each other than they are to CG2941. In contrast, CG2941 shares more sequences with the orthologous sequences in other more distantly related Drosophilids than do CG32786 or CG32783. Thus, we infer that CG2941 is ancestral to the CG32786 and CG32783 gene pair.

**Figure 6.**
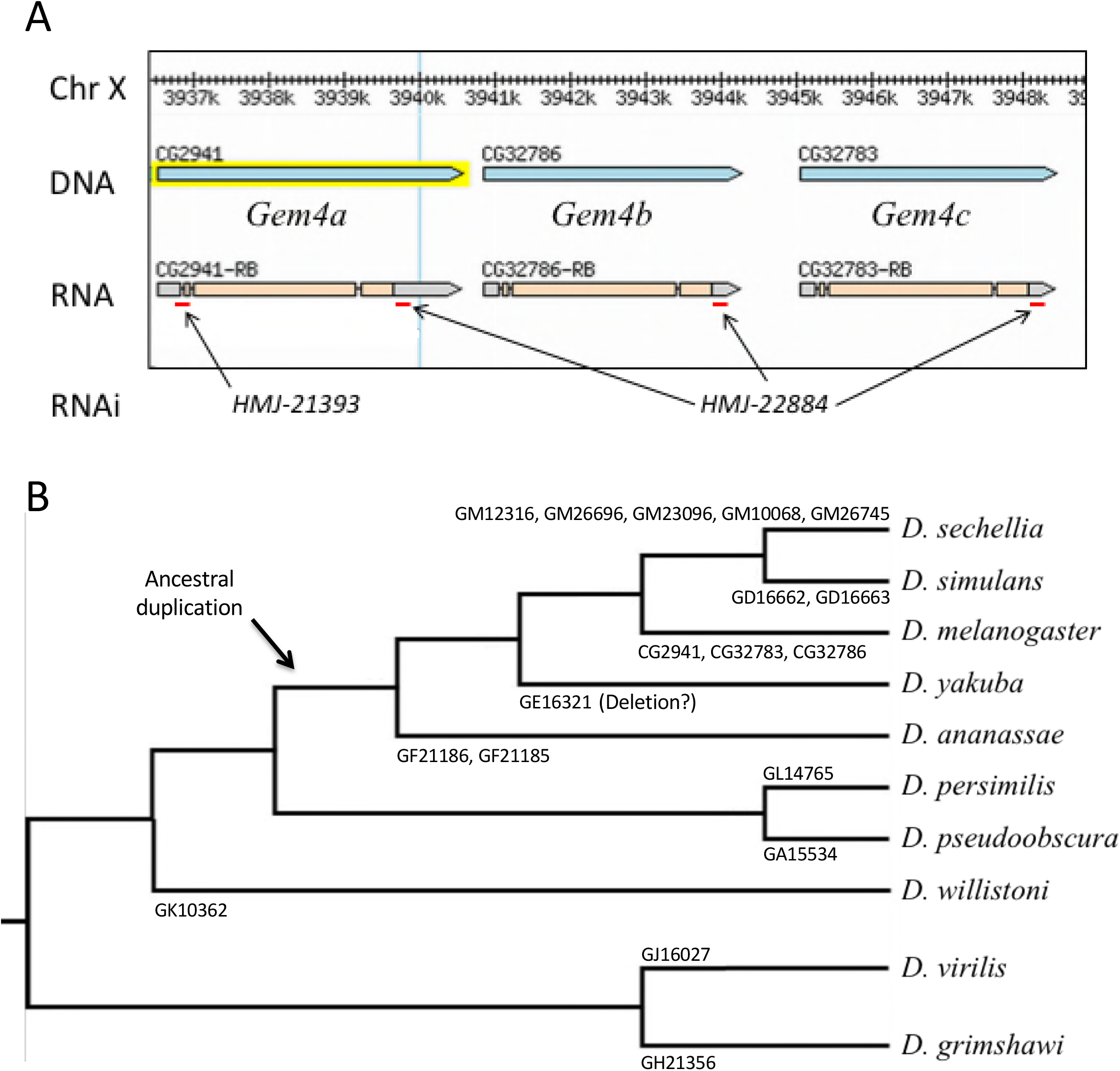
Organization and phylogeny of Gemin4/CG2941-like genes in Drosophilids. (A) Browser shot of *D. melanogaster* CG2941 paralogs, showing their relative location and exon-intron structure. Red dashes indicate regions targeted by short hairpin (sh)RNAs expressed from Gal4-inducible RNAi constructs, see text for details. (B) *Drosophila* phylogenetic tree showing gene name and number of respective Gemin4 orthologs within each lineage. Following their divergence from the obscura group (*D. pseudoobscura* and *D. persimilis*), potential duplications and deletions within the melanogaster group (*D. sechellia, D. melanogaster, D. simulans, D. yakuba, and D. annanasae*) are illustrated.

Interestingly, this region of the genome appears to be somewhat fluid, particularly within the melanogaster group, which includes *D. sechellia, D. melanogaster, D. simulans, D. yakuba, and D. annanasae*. Phylogenetic analysis of the number of CG2941-like genes in various Drosophilid genomes (Fig. 6B) suggests that an ancestral duplication of CG2941 occurred sometime between the divergence of the melanogaster group and the obscura group, the latter of which contains *D. pseudoobscura* and *D. persimilis*. The hawaiian (represented by *D. grimshawi*) and virilis (*D. virilis*) groups each only have one CG2941-like gene and thus serve as outgroups for this analysis. Within the melanogaster group there appears to be ongoing genetic rearrangement of this genomic region, as certain species have up to five different copies of this gene, whereas others have only a single copy (Fig. 6B). For ease of future identification, we suggest the following nomenclature for the *D. melanogaster* genes: *Gem4a (CG2941/Gaulos), Gem4b (CG32786) and Gem4c (CG32783)*.

### Gemin4 gene function is essential for viability in Drosophila

The FlyAtlas anatomical and developmental expression database (Chintapalli *et al.* 2007; Leader *et al.* 2018) shows that *CG2941* is expressed ubiquitously, albeit at relatively low levels. Its highest levels of expression are in the larval central nervous system and the adult ovary. Because the sequences of the three *Gem4* paralogs are so similar, the function and expression levels of the other two genes is unclear. Lanfranco et al. (2017) employed an RNA interference transgene targeting *CG2941*, but did not mention the existence of the other two paralogous genes in their publication. We note that this transgene (VDRC 52356), obtained from the Vienna Drosophila Resource Center, expresses a 339bp dsRNA that targets all three *Gem4* paralogs (*CG2941, CG32786* and *CG32783*). Furthermore, the deficiency, *Df(1)ED6716*, used in the original genetic interaction screen also uncovers all three paralogs.

To confirm and extend these studies, we sought to determine whether or not loss of *CG2941* might be compensated by the presence of the other paralogs. We therefore carried out RNAi using two different shRNA expressing TRiP lines obtained from the Transgenic RNAi Project (Perkins *et al.* 2015). As shown in Fig. 6A, HMJ-21393 specifically targets a region near the 5’-end of the *Gem4a/CG2941* transcript, whereas the HMJ-22884 construct targets a 3’-UTR sequence that is shared by all three transcripts.

In Fig. 7A, we used the Gal4/UAS system to drive these two UAS-shRNA constructs ubiquitously using either *daughterless*-Gal4 (Da-Gal4) or *tubulin*-Gal4 (Tub-Gal4). Expression of either drivers or the responders alone had little to no effect on organismal viability (Fig. 7A). However, ubiquitous expression of the shRNA that targets all three genes (HMJ-22884) was essentially larval lethal (Fig. 7A). The phenotype of animals expressing the HMJ-21933 construct that specifically targets *CG2941* was slightly less severe, as most of the animals complete larval development (Da-Gal4 x HMJ-21933), and roughly 20% of them eclose as adults. The phenotype of the Tub-Gal4 x HMJ-21933 cross was even more severe, as only ∼30% of the animals reached pupal stages and none progressed to adulthood (Fig. 7A). These findings are consistent with those in mice (Meier *et al.* 2018), showing that *Gem4* is an essential gene. The data also suggest that *Gem4b/CG32786* and *Gem4c/CG32783* can partially compensate for loss of *Gem4a/CG2941.* However, in the absence of specific genetic lesions and complementation analysis, it is difficult to make firm conclusions.

**Figure 7.**
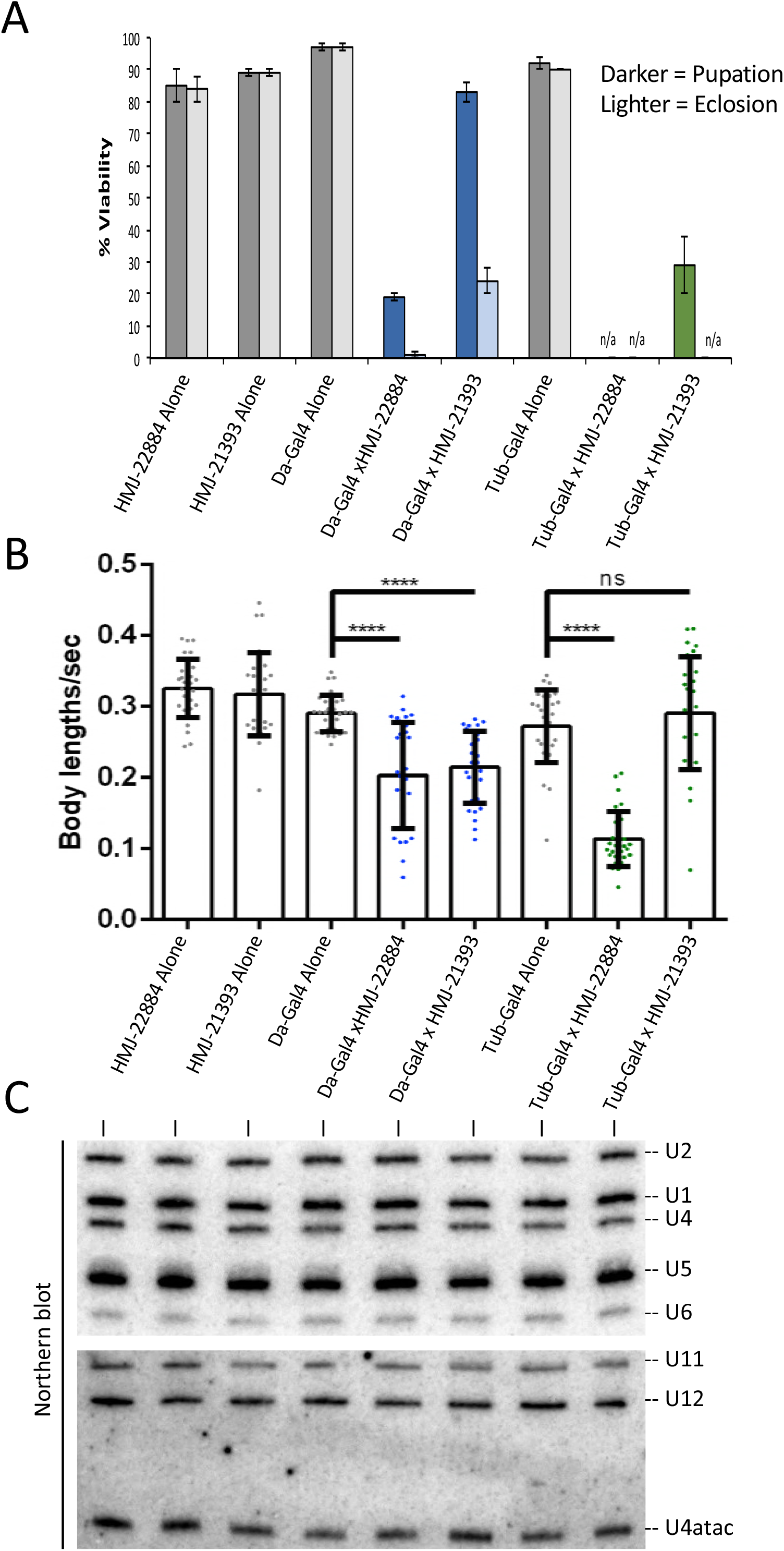
*CG2941* and its paralogs are essential for viability and locomotion independent of snRNP biogenesis. (**A**) *daughterless* (Da) or *tubulin* (Tub) Gal4 drivers were crossed to RNAi lines specifically targeting *CG2941* (HMJ-21393) or all three paralogs (HMJ-22884). 100 larvae per cross were selected and viability was measured based on the number of larvae that pupated (darker bars) and that eclosed (lighter bars). “n/a” represents a category in which there were no pupae (pupation) or adults (eclosion) to count. Error bars represent standard error. (**B**) Wandering third instar larvae from the same crosses in (A) were recorded and locomotor behavior was measured using the wrMTrck plug-in for Fiji/ImageJ. (**C**) Early third instar larvae (73-77 hours post-egg laying) from the same crosses in (A) were collected and RNA was extracted to measure snRNA levels via northern blotting. Probe sequences are described in Materials and Methods.

To investigate the consequences of Gem4 loss of function, we carried out larval locomotion assays and northern blotting analysis. Wandering third instar larvae were used to record their movement on a molasses agar plate. The videos were then converted and analyzed using the wrmTrck plug-in of Fiji/ImageJ to generate a measurement of body lengths per second (BLPS), which takes into account the speed and size of each larva. For northern blotting, total RNA was harvested from early third instar (72-76hr) larvae, just prior to the beginning of the lethal phase of the RNAi. As shown in Fig. 7C, we did not detect signficant reductions in the levels of either the major (U1, U2, U4 and U5) or the minor (U11, U12 and U4atac) Sm-class snRNAs. Due to the long half-lives of spliceosomal snRNPs in cultured mammalian cells (1-3 days, depending on the snRNA; Sauterer *et al.* 1988), this finding is perhaps not so surprising. Thus the presumptive loss of Gem4 function in snRNP biogenesis may not have had time to affect these animals. Given that complete loss of SMN protein only results in a ∼50-60% reduction of U1-U5 snRNAs at this stage of development (Garcia *et al.* 2016), it is also unsurprising that knockdown of Gem4 has a less dramatic effect. We conclude that the larval lethality associated with *Gem4* loss of function is not due to a concomitant loss of snRNPs.

### Conservation of Gemin4 among Dipteran genomes

In the ten years since Kroiss et al. (2008) first suggested that Gem4 was missing from the Dipteran SMN complex, there have been numerous hints to the contrary. As early as 2009, raw mass spectrometry data released by the DPiM (*Drosophila* Protein interaction Map) project showed that CG2941 co-precipitates with ectopically expressed, epitope-tagged SMN in S2 cells (https://interfly.med.harvard.edu). Subsequent quality control steps apparently removed CG2941 from the list of potential SMN interactors despite the fact that it also co-purifies with epitope tagged Gem2 and Gem3 (Guruharsha *et al.* 2011; Guruharsha *et al.* 2012). On the basis of biochemical purifications from fly embryos and S2 cells, we and others have speculated that CG2941 might well be a *bona fide* core member of the SMN complex (Sen *et al.* 2013; Gray *et al.* 2016). However, in the absence of additional genetic, phylogenetic and biochemical evidence linking endogenous CG2941 to SMN, the conservation of Gem4 remained an open question.

Three new lines of experimentation demonstrate that *CG2941* is indeed *Gem4*. First, Lanfranco et al. (2017) found that an N-terminally truncated *Gem3^∆^N* construct interacts genetically with a deficiency that uncovers *CG2941/Gem4a, CG32786/Gem4b* and *CG32783/Gem4c*. They also showed that RNAi-mediated knockdown of all three genes enhanced the phenotype of this dominant negative *Gem3^∆^N* transgene. Second, ongoing metazoan genome sequencing efforts allowed us to more confidently predict *Gem4* orthologs on the basis of primary sequence (Fig. S1 and data not shown). Third, we found that endogenous CG2941, CG32786 and CG32783 co-purify with SMN expressed from its native promoter *in vivo* in fly embryos (Fig. 5). The relative enrichment and number of peptides corresponding to CG2941 in the mass spectrometry experiment was similar to that of the other Gemins (Fig. 5A). These findings lead us to conclude that *Gemin4* has been retained in the genomes of *Drosophila* and other dipterans.

### SMN and the evolution of Gemin subcomplexes

The human SMN complex can be subdivided into several distinct subunits (Battle *et al.* 2007; Otter *et al.* 2007). SMN and Gem2 form an oligomeric heterodimer (SMN•Gem2)_n_ that makes up the core of the complex (Fischer *et al.* 1997; Gupta *et al.* 2015). Gem3 binds directly and independently to Gem4, tethering them both to SMN•Gem2 (Charroux *et al.* 1999; Charroux *et al.* 2000). Oligomerization of SMN appears to be required for Gem3 to enter the complex (Praveen *et al.* 2014). Gem5 is a large (175 kD) WD-repeat protein that recruits RNA substrates to the SMN complex (Battle *et al.* 2006b) via subdomains that bind to the m7G-cap and Sm-site, respectively (Xu *et al.* 2016). Thus Gem5 can be viewed as an RNP subunit of the SMN complex. Finally, Gem6 and Gem7 heterodimerize (Baccon *et al.* 2002; Ma *et al.* 2005) and recruit Gem8 (Carissimi *et al.* 2006) to form a Gem6-7-8 subunit, the function of which is unknown. As shown in Figs. 2 and 3, these three proteins appear to have been lost from Drosophilids, but retained in the genomes of other insects.

An important question raised by our findings is whether or not the functions normally carried out by Gem6, Gem7 and Gem8 may have been taken over by CG14164 (Hez/Lsm12b), CG31950 (Sbat/Naa38) and CG2371 (Vlet/CommD10). Lanfranco et al. (2017) showed that these three proteins are each capable of forming complexes with exogenously expressed SMN when they are transfected into S2 cells. And given the opportunity to interact in a directed yeast two-hybrid screen, Sbat/Naa38 scored positively for interaction with Gem3, Hez/Lsm12b and Vlet/CommD10 (Lanfranco *et al.* 2017). Interestingly, both Hez and Sbat are predicted contain an Sm-fold, also known as a small beta barrel (Youkharibache *et al.* 2018). This structure is characterized by five short beta strands that form a closed domain wherein the first strand is hydrogen bonded to the last (Arluison *et al.* 2006).

Small beta barrel containing proteins exhibit a strong tendency to form higher-order structures, as exemplified by the Sm and Lsm proteins, found in all three domains of life (Youkharibache *et al.* 2018). Thus, despite the fact that Hez, Sbat and Vlet are not orthologous to Gem6, Gem7 and Gem8, respectively, it remains formally possible that they have been evolutionarily co-opted into the SMN complex in flies. Our inability to identify these three proteins as endogenous SMN binding partners by mass spectrometry (Fig. 5) argues against this idea. However, stable protein interactions are not required to elicit important biological outcomes, so additional experiments will be needed to conclusively demonstrate a role for Hez/Lsm12b, Sbat/Naa38 and Vlet/CommD10 in SMN biology.

### Considerations and Prospects

In the mean time, several interesting possibilities suggest themselves for future investigation. We and others have hypothesized that the SMN complex may function as a hub for various cellular signaling pathways, in addition to its role in chaperoning snRNP biogenesis (Raimer *et al.* 2017; Gray *et al.* 2018 and references therein). As shown in Fig. 4, the fruitfly CommD10 orthologue is (Vlet/CG2371). Intriguingly, human CommD (Copper metabolism Murr1 domain) proteins can form homo- and hetero-dimers (Burstein *et al.* 2005) and are involved in a variety of cellular pathways including endosomal membrane trafficking and the inhibition of NF-kB signaling (Bartuzi *et al.* 2013; Mallam and Marcotte 2017). Both SMN (Kim and Choi 2017) and Gem3 (Shin *et al.* 2014) have been implicated in NF-kB related pathways. It is tempting to speculate that human Gem8 might play a role in linking the SMN complex to NF-kB signaling by interacting with, or otherwise functioning as, a CommD-like protein. Given the potential for Sbat/Naa38 to interact with Gem3 (Lanfranco *et al.* 2017), perhaps the Gem6-7-8 subcomplex functions as a regulatory subunit that modulates the activity of SMN and/or Gem3.

Irrespective of any putative role in cellular signaling, the fact that Sbat/Naa38 contains an Sm-fold may help to explain several interesting observations in the literature. Metazoan Naa38 (a.k.a. Lsmd1, Mak31) is an auxiliary subunit of NatC (N-terminal acetyltransferase C; Starheim *et al.* 2009). N(alpha)-acetyltransferases are enzymes that consist of a catalytic subunit and one or two auxiliary subunits (Aksnes *et al.* 2016). The auxiliary subunits modulate the activity and substrate specificity of the catalytic subunit. Furthermore, they mediate co-translational binding to the 60S ribosome, in a region that is located near the nascent polypeptide exit tunnel (reviewed in Aksnes *et al.* 2016). This latter point merits attention for two reasons.

First, Fischer and colleagues recently reported data suggesting that, following their translation, Sm proteins can remain bound to the ribosome near the exit tunnel, dissociating only after binding to the assembly chaperone pICln (Paknia *et al.* 2016). These authors hypothesize the existence of a quality control hub for chaperone-mediated protein assembly, located on the ribosome. Whether or not Sm protein heterodimers (e.g. Lsm10/11 and SmD1/D2) actually bind to the nascent peptide tunnel region of the ribosome *in viv*o is unclear. However, the fact that metazoan Naa38 is structurally similar to Sm proteins provides a plausible mechanism for their binding to the ribosome immediately following translation. Given that Gem6 and Gem7 are also members of the Sm-like superfamily of proteins (Ma *et al.* 2005) it is conceivable that Sbat/Naa38 could dimerize with other Sm-like proteins (e.g. Hez/Lsm12b) in *Drosophila*.

Second, Naa38 might not be a *bona fide* member of the SMN complex (in flies or any other species), but it could potentially interact with SMN as part of its *canonical* function in N-terminal protein acetylation. More than 80% of human proteins are cotranslationally modified on their N-termini (Arnesen *et al.* 2009), however the functional impact of this modification is largely unknown. Most proteins do not retain their N-terminal Met residue, and its removal by methionine aminopeptidases frequently leads to acetylation of the resulting N-terminus, particularly if the second residue is Ala, Val, Ser, Thr or Cys (Hwang *et al.* 2010). Interestingly, the N-terminal Ala2 residue of SMN is known to be acetylated in human cells (Van Damme *et al.* 2012), and an A2G missense mutation in the *SMN1* gene is known to cause a mild form of SMA when *SMN2* is present in a single copy (Parsons *et al.* 1998). This mutation is puzzling because, with the exception of Ala2, the N-terminal 15aa of SMN (i.e. upstream of the Gemin2 binding domain) are very poorly conserved. Moreover, changing Ala2 to Gly is predicted to reduce the probability of N-terminal acetylation and recognition by the N-end-rule proteasomal degradation pathway (Hwang *et al.* 2010). These findings suggest that the phenotype of the A2G mutation in humans is due to loss of N-terminal acetylation of SMN.

In conclusion, it seems unlikely that three different proteins (CG14164, CG31950, CG2371) derived from three different biological contexts might be co-opted into a novel Gemin subcomplex. A loss of the Gem6-7-8 subunit from the SMN complex in flies would suggest that either this subunit is not essential for basal metazoan viability or that other factors have compensated for deficiency of these proteins in *Drosophila*. Additional experiments will be needed to rule in, or rule out, any such functional adaptation. In contrast, the identification of Gem4 via PSI-BLAST in a variety of different insect genomes including the Diptera, Hemiptera, Lepidoptera, and Hymenoptera (this work) indicates that this protein is widely conserved. Moreover, genetic loss of function studies (Fig. 7; Lanfranco *et al.* 2017; Meier *et al.* 2018) strongly suggest that Gem4 is essential for metazoan viability.

## Figure Legends

**Figure S1.**
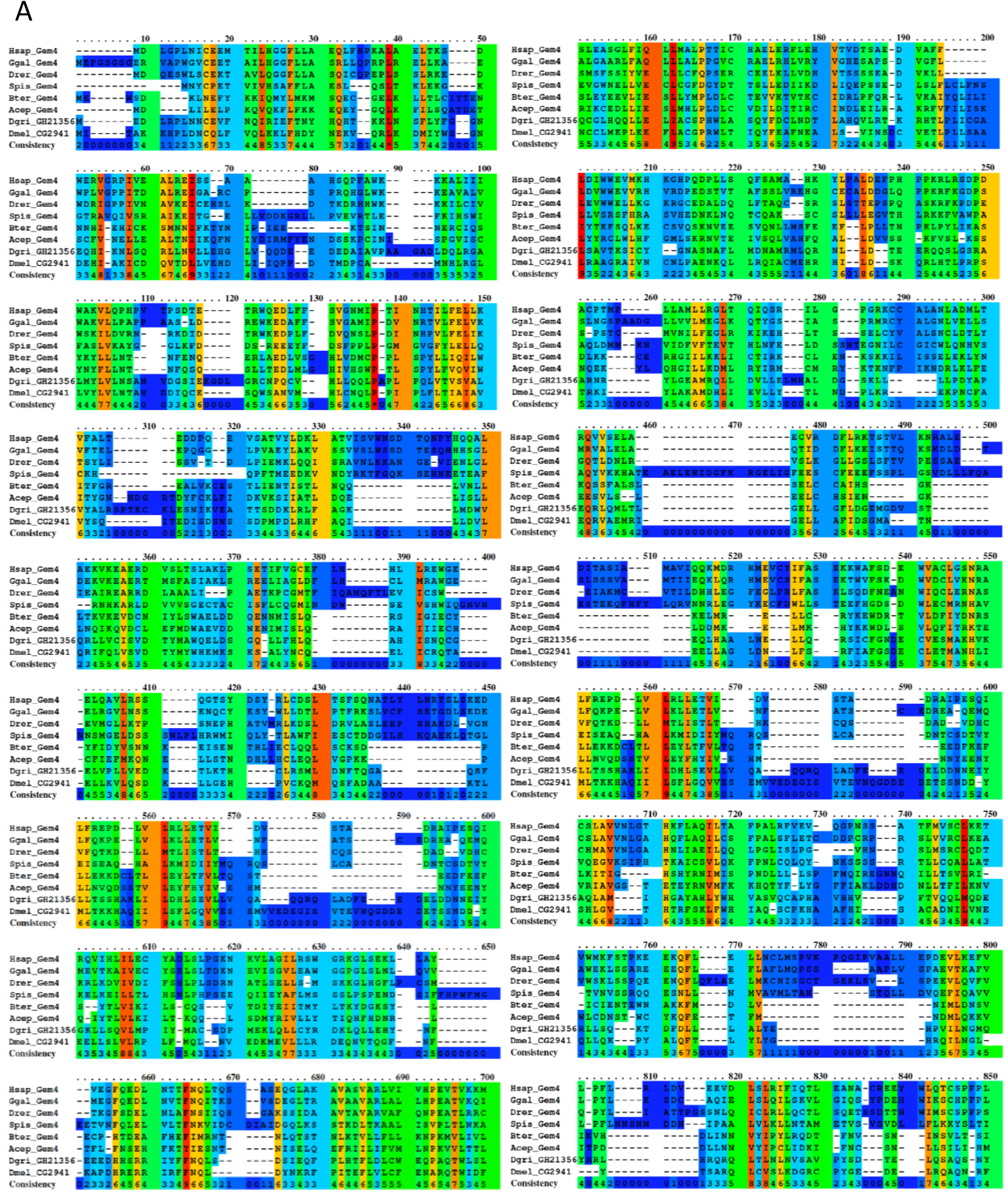
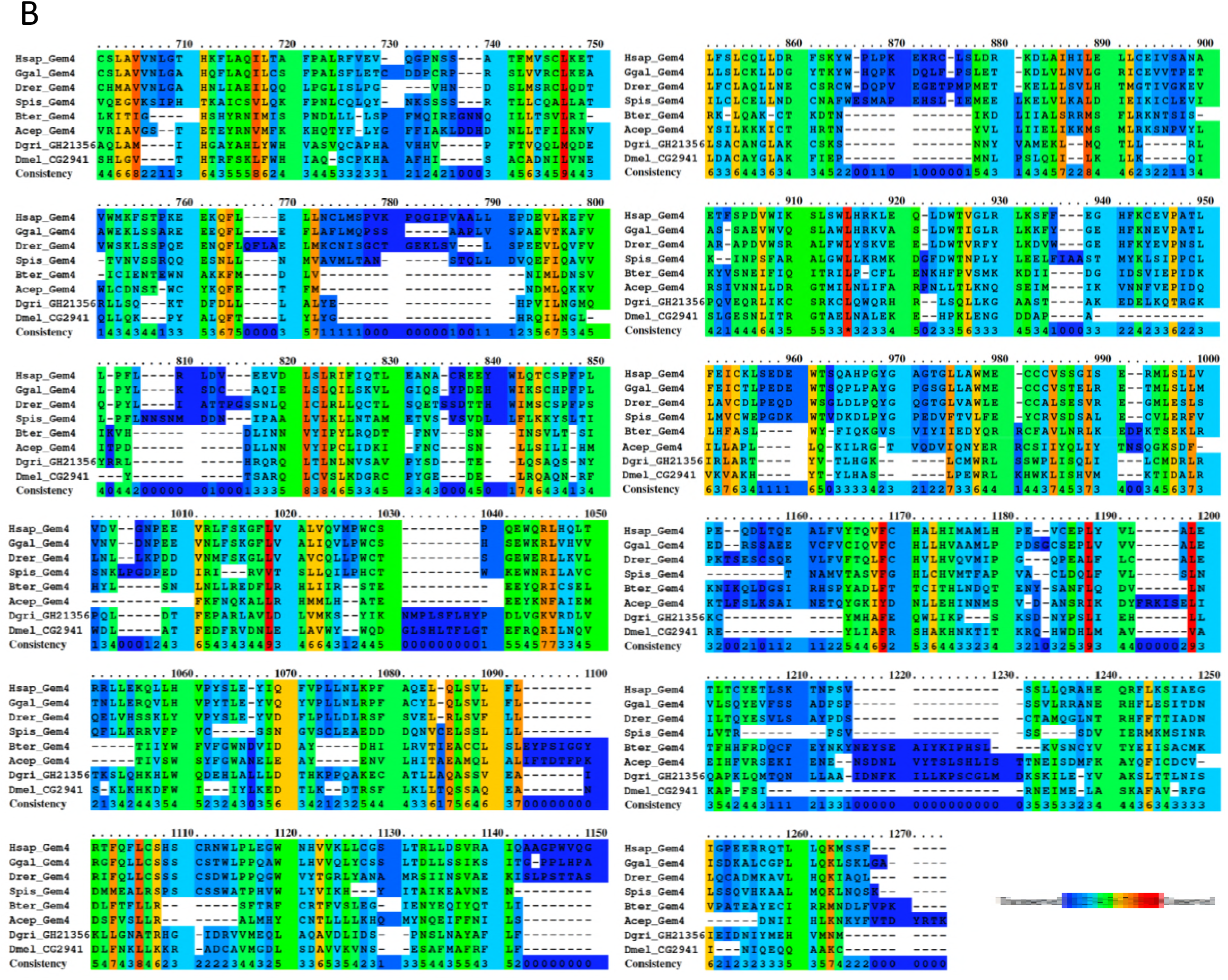
Amino acid alignment of Gemin4 orthologs from a variety of species, including: *Homo sapiens* (Hsap), *Gallus gallus* (Ggal) *Danio rerio* (Drer), *Stylophora pistillata* (Spis), *Bombus terrestris* (Bter), *Atta cephalotes* (Acep), *Drosophila grimshawi* (Dgri), and *Drosophila melanogaster* (Dmel). (**A**) Alignment of Dmel_CG2941 with residues 1-850 of metazoan Gemin4. (**B**) Alignment of Dmel_CG2941 with Gemin4 residues 700-1274. (A-B) The degree of conservation in the sequence is indicated by a coloration spectrum running from blue to red, with blue indicating regions of low conservation and red highlighting those that are highly conserved.

## References

Aksnes, H., A. Drazic, M. Marie and T. Arnesen, 2016 First Things First: Vital Protein Marks by N-Terminal Acetyltransferases. Trends Biochem Sci 41: 746-760.

Albrecht, M., and T. Lengauer, 2004 Novel Sm-like proteins with long C-terminal tails and associated methyltransferases. FEBS Lett 569: 18-26.

Altschul, S. F., T. L. Madden, A. A. Schaffer, J. Zhang, Z. Zhang et al., 1997 Gapped BLAST and PSI-BLAST: a new generation of protein database search programs. Nucleic Acids Res 25: 3389-3402.

Arluison, V., C. Mura, M. R. Guzman, J. Liquier, O. Pellegrini et al., 2006 Three-dimensional structures of fibrillar Sm proteins: Hfq and other Sm-like proteins. J Mol Biol 356: 86-96.

Arnesen, T., P. Van Damme, B. Polevoda, K. Helsens, R. Evjenth et al., 2009 Proteomics analyses reveal the evolutionary conservation and divergence of N-terminal acetyltransferases from yeast and humans. Proc Natl Acad Sci U S A 106: 8157-8162.

Baccon, J., L. Pellizzoni, J. Rappsilber, M. Mann and G. Dreyfuss, 2002 Identification and characterization of Gemin7, a novel component of the survival of motor neuron complex. J Biol Chem 277: 31957-31962.

Bartuzi, P., M. H. Hofker and B. van de Sluis, 2013 Tuning NF-kappaB activity: a touch of COMMD proteins. Biochim Biophys Acta 1832: 2315-2321.

Battle, D. J., M. Kasim, J. Wang and G. Dreyfuss, 2007 SMN-independent subunits of the SMN complex. Identification of a small nuclear ribonucleoprotein assembly intermediate. J Biol Chem 282: 27953-27959.

Battle, D. J., M. Kasim, J. Yong, F. Lotti, C. K. Lau et al., 2006a The SMN complex: an assembly machine for RNPs. Cold Spring Harb Symp Quant Biol 71: 313-320.

Battle, D. J., C. K. Lau, L. Wan, H. Deng, F. Lotti et al., 2006b The Gemin5 protein of the SMN complex identifies snRNAs. Mol Cell 23: 273-279.

Borg, R., and R. J. Cauchi, 2013 The Gemin associates of survival motor neuron are required for motor function in Drosophila. PLoS One 8: e83878.

Borg, R. M., R. Bordonne, N. Vassallo and R. J. Cauchi, 2015 Genetic Interactions between the Members of the SMN-Gemins Complex in Drosophila. PLoS One 10: e0130974.

Bradrick, S. S., and M. Gromeier, 2009 Identification of gemin5 as a novel 7-methylguanosine cap-binding protein. PLoS One 4: e7030.

Burghes, A. H., and C. E. Beattie, 2009 Spinal muscular atrophy: why do low levels of survival motor neuron protein make motor neurons sick? Nat Rev Neurosci 10: 597-609.

Burstein, E., J. E. Hoberg, A. S. Wilkinson, J. M. Rumble, R. A. Csomos et al., 2005 COMMD proteins, a novel family of structural and functional homologs of MURR1. J Biol Chem 280: 22222-22232.

Carissimi, C., J. Baccon, M. Straccia, P. Chiarella, A. Maiolica et al., 2005 Unrip is a component of SMN complexes active in snRNP assembly. FEBS Lett 579: 2348-2354.

Carissimi, C., L. Saieva, J. Baccon, P. Chiarella, A. Maiolica et al., 2006 Gemin8 is a novel component of the survival motor neuron complex and functions in small nuclear ribonucleoprotein assembly. J Biol Chem 281: 8126-8134.

Cauchi, R. J., L. Sanchez-Pulido and J. L. Liu, 2010 Drosophila SMN complex proteins Gemin2, Gemin3, and Gemin5 are components of U bodies. Exp Cell Res 316: 2354-2364.

Charroux, B., L. Pellizzoni, R. A. Perkinson, A. Shevchenko, M. Mann et al., 1999 Gemin3: A novel DEAD box protein that interacts with SMN, the spinal muscular atrophy gene product, and is a component of gems. J Cell Biol 147: 1181-1194.

Charroux, B., L. Pellizzoni, R. A. Perkinson, J. Yong, A. Shevchenko et al., 2000 Gemin4. A novel component of the SMN complex that is found in both gems and nucleoli. J Cell Biol 148: 1177-1186.

Chaytow, H., Y. T. Huang, T. H. Gillingwater and K. M. E. Faller, 2018 The role of survival motor neuron protein (SMN) in protein homeostasis. Cell Mol Life Sci.

Chintapalli, V. R., J. Wang and J. A. Dow, 2007 Using FlyAtlas to identify better Drosophila melanogaster models of human disease. Nat Genet 39: 715-720.

Coady, T. H., and C. L. Lorson, 2011 SMN in spinal muscular atrophy and snRNP biogenesis. Wiley Interdiscip Rev RNA 2: 546-564.

Curmi, F., and R. J. Cauchi, 2018 The multiple lives of DEAD-box RNA helicase DP103/DDX20/Gemin3. Biochem Soc Trans 46: 329-341.

Di, Y., J. Li, Y. Zhang, X. He, H. Lu et al., 2003 HCC-associated protein HCAP1, a variant of GEMIN4, interacts with zinc-finger proteins. J Biochem 133: 713-718.

Fallini, C., G. J. Bassell and W. Rossoll, 2012 Spinal muscular atrophy: the role of SMN in axonal mRNA regulation. Brain Res 1462: 81-92.

Fischer, U., C. Englbrecht and A. Chari, 2011 Biogenesis of spliceosomal small nuclear ribonucleoproteins. Wiley Interdiscip Rev RNA 2: 718-731.

Fischer, U., Q. Liu and G. Dreyfuss, 1997 The SMN-SIP1 complex has an essential role in spliceosomal snRNP biogenesis. Cell 90: 1023-1029.

Garcia, E. L., Y. Wen, K. Praveen and A. G. Matera, 2016 Transcriptomic comparison of Drosophila snRNP biogenesis mutants reveals mutant-specific changes in premRNA processing: implications for spinal muscular atrophy. RNA 22: 1215-1227.

Gates, J., G. Lam, J. A. Ortiz, R. Losson and C. S. Thummel, 2004 rigor mortis encodes a novel nuclear receptor interacting protein required for ecdysone signaling during Drosophila larval development. Development 131: 25-36.

Gray, K. M., K. A. Kaifer, D. Baillat, Y. Wen, T. R. Bonacci et al., 2018 Self-oligomerization regulates stability of survival motor neuron protein isoforms by sequestering an SCF(Slmb) degron. Mol Biol Cell 29: 96-110.

Gray, K. M., Y. Wen and A. G. Matera, 2016 A novel component of the Drosophila SMN complex? CureSMA Proc. 20: 55.

Grimmler, M., S. Otter, C. Peter, F. Muller, A. Chari et al., 2005 Unrip, a factor implicated in cap-independent translation, associates with the cytosolic SMN complex and influences its intracellular localization. Hum Mol Genet 14: 3099-3111.

Groen, E. J. N., K. Talbot and T. H. Gillingwater, 2018 Advances in therapy for spinal muscular atrophy: promises and challenges. Nat Rev Neurol 14: 214-224.

Grundhoff, A. T., E. Kremmer, O. Tureci, A. Glieden, C. Gindorf et al., 1999 Characterization of DP103, a novel DEAD box protein that binds to the Epstein-Barr virus nuclear proteins EBNA2 and EBNA3C. J Biol Chem 274: 19136-19144.

Gupta, K., R. Martin, R. Sharp, K. L. Sarachan, N. S. Ninan et al., 2015 Oligomeric Properties of Survival Motor Neuron.Gemin2 Complexes. J Biol Chem 290: 20185-20199.

Guruharsha, K. G., R. A. Obar, J. Mintseris, K. Aishwarya, R. T. Krishnan et al., 2012 Drosophila protein interaction map (DPiM): a paradigm for metazoan protein complex interactions. Fly (Austin) 6: 246-253.

Guruharsha, K. G., J. F. Rual, B. Zhai, J. Mintseris, P. Vaidya et al., 2011 A protein complex network of Drosophila melanogaster. Cell 147: 690-703.

Hamilton, G., and T. H. Gillingwater, 2013 Spinal muscular atrophy: going beyond the motor neuron. Trends Mol Med 19: 40-50.

Hunt, S. L., J. J. Hsuan, N. Totty and R. J. Jackson, 1999 unr, a cellular cytoplasmic RNA-binding protein with five cold-shock domains, is required for internal initiation of translation of human rhinovirus RNA. Genes Dev 13: 437-448.

Hutvágner, G., and P. D. Zamore, 2002 A microRNA in a multiple-turnover RNAi enzyme complex. Science 297: 2056-2060.

Hwang, C. S., A. Shemorry and A. Varshavsky, 2010 N-terminal acetylation of cellular proteins creates specific degradation signals. Science 327: 973-977.

Jablonka, S., B. Holtmann, G. Meister, M. Bandilla, W. Rossoll et al., 2002 Gene targeting of Gemin2 in mice reveals a correlation between defects in the biogenesis of U snRNPs and motoneuron cell death. Proc Natl Acad Sci U S A 99: 10126-10131.

Jin, W., Y. Wang, C. P. Liu, N. Yang, M. Jin et al., 2016 Structural basis for snRNA recognition by the double-WD40 repeat domain of Gemin5. Genes Dev 30: 2391-2403.

Khokhar, A., N. Chen, J. P. Yuan, Y. Li, G. N. Landis et al., 2008 Conditional switches for extracellular matrix patterning in Drosophila melanogaster. Genetics 178: 1283-1293.

Kim, E. K., and E. J. Choi, 2017 SMN1 functions as a novel inhibitor for TRAF6-mediated NF-kappaB signaling. Biochim Biophys Acta Mol Cell Res 1864: 760-770.

Kim, E. K., K. T. Noh, J. H. Yoon, J. H. Cho, K. W. Yoon et al., 2007 Positive regulation of ASK1-mediated c-Jun NH(2)-terminal kinase signaling pathway by the WD-repeat protein Gemin5. Cell Death Differ 14: 1518-1528.

Kroiss, M., J. Schultz, J. Wiesner, A. Chari, A. Sickmann et al., 2008 Evolution of an RNP assembly system: a minimal SMN complex facilitates formation of UsnRNPs in Drosophila melanogaster. Proc Natl Acad Sci U S A 105: 10045-10050.

Lanfranco, M., R. Cacciottolo, R. M. Borg, N. Vassallo, F. Juge et al., 2017 Novel interactors of the Drosophila Survival Motor Neuron (SMN) Complex suggest its full conservation. FEBS Lett 591: 3600-3614.

Lau, C. K., J. L. Bachorik and G. Dreyfuss, 2009 Gemin5-snRNA interaction reveals an RNA binding function for WD repeat domains. Nat Struct Mol Biol 16: 486-491.

Leader, D. P., S. A. Krause, A. Pandit, S. A. Davies and J. A. T. Dow, 2018 FlyAtlas 2: a new version of the Drosophila melanogaster expression atlas with RNA-Seq, miRNA-Seq and sex-specific data. Nucleic Acids Res 46: D809-D815.

Lee, J., E. Yoo, H. Lee, K. Park, J. H. Hur et al., 2017 LSM12 and ME31B/DDX6 Define Distinct Modes of Posttranscriptional Regulation by ATAXIN-2 Protein Complex in Drosophila Circadian Pacemaker Neurons. Mol Cell 66: 129-140 e127.

Lefebvre, S., L. Burglen, S. Reboullet, O. Clermont, P. Burlet et al., 1995 Identification and characterization of a spinal muscular atrophy-determining gene. Cell 80: 155-165.

Liu, Q., U. Fischer, F. Wang and G. Dreyfuss, 1997 The spinal muscular atrophy disease gene product, SMN, and its associated protein SIP1 are in a complex with spliceosomal snRNP proteins. Cell 90: 1013-1021.

Lorson, C. L., J. Strasswimmer, J. M. Yao, J. D. Baleja, E. Hahnen et al., 1998 SMN oligomerization defect correlates with spinal muscular atrophy severity. Nat Genet 19: 63-66.

Ma, Y., J. Dostie, G. Dreyfuss and G. D. Van Duyne, 2005 The Gemin6-Gemin7 heterodimer from the survival of motor neurons complex has an Sm protein-like structure. Structure 13: 883-892.

Maine, G. N., and E. Burstein, 2007 COMMD proteins: COMMing to the scene. Cell Mol Life Sci 64: 1997-2005.

Mallam, A. L., and E. M. Marcotte, 2017 Systems-wide Studies Uncover Commander, a Multiprotein Complex Essential to Human Development. Cell Syst 4: 483-494.

Martin, R., K. Gupta, N. S. Ninan, K. Perry and G. D. Van Duyne, 2012 The survival motor neuron protein forms soluble glycine zipper oligomers. Structure 20: 1929-1939.

Matera, A. G., R. M. Terns and M. P. Terns, 2007 Non-coding RNAs: lessons from the small nuclear and small nucleolar RNAs. Nat Rev Mol Cell Biol 8: 209-220.

Matera, A. G., and Z. Wang, 2014 A day in the life of the spliceosome. Nat Rev Mol Cell Biol 15: 108-121.

Meier, I. D., M. P. Walker and A. G. Matera, 2018 Gemin4 is an essential gene in mice, and its overexpression in human cells causes relocalization of the SMN complex to the nucleoplasm. Biol Open 7: bio032409.

Meister, G., D. Buhler, B. Laggerbauer, M. Zobawa, F. Lottspeich et al., 2000 Characterization of a nuclear 20S complex containing the survival of motor neurons (SMN) protein and a specific subset of spliceosomal Sm proteins. Hum Mol Genet 9: 1977-1986.

Meister, G., M. Landthaler, L. Peters, P. Y. Chen, H. Urlaub et al., 2005 Identification of novel argonaute-associated proteins. Curr Biol 15: 2149-2155.

Mouillet, J. F., X. Yan, Q. Ou, L. Jin, L. J. Muglia et al., 2008 DEAD-box protein-103 (DP103, Ddx20) is essential for early embryonic development and modulates ovarian morphology and function. Endocrinology 149: 2168-2175.

Mourelatos, Z., J. Dostie, S. Paushkin, A. Sharma, B. Charroux et al., 2002 miRNPs: a novel class of ribonucleoproteins containing numerous microRNAs. Genes Dev 16: 720-728.

Narayanan, U., T. Achsel, R. Luhrmann and A. G. Matera, 2004 Coupled in vitro import of U snRNPs and SMN, the spinal muscular atrophy protein. Mol Cell 16: 223-234.

Nash, L. A., J. K. Burns, J. W. Chardon, R. Kothary and R. J. Parks, 2016 Spinal Muscular Atrophy: More than a Disease of Motor Neurons? Curr Mol Med 16: 779-792.

Otter, S., M. Grimmler, N. Neuenkirchen, A. Chari, A. Sickmann et al., 2007 A comprehensive interaction map of the human survival of motor neuron (SMN) complex. J Biol Chem 282: 5825-5833.

Paknia, E., A. Chari, H. Stark and U. Fischer, 2016 The Ribosome Cooperates with the Assembly Chaperone pICln to Initiate Formation of snRNPs. Cell Rep 16: 3103-3112.

Parsons, D. W., P. E. McAndrew, S. T. Iannaccone, J. R. Mendell, A. H. Burghes et al., 1998 Intragenic telSMN mutations: frequency, distribution, evidence of a founder effect, and modification of the spinal muscular atrophy phenotype by cenSMN copy number. Am J Hum Genet 63: 1712-1723.

Paushkin, S., A. K. Gubitz, S. Massenet and G. Dreyfuss, 2002 The SMN complex, an assemblyosome of ribonucleoproteins. Curr Opin Cell Biol 14: 305-312.

Pellizzoni, L., B. Charroux and G. Dreyfuss, 1999 SMN mutants of spinal muscular atrophy patients are defective in binding to snRNP proteins. Proc Natl Acad Sci U S A 96: 11167-11172.

Perkins, L. A., L. Holderbaum, R. Tao, Y. Hu, R. Sopko et al., 2015 The Transgenic RNAi Project at Harvard Medical School: Resources and Validation. Genetics 201: 843-852.

Pineiro, D., J. Fernandez-Chamorro, R. Francisco-Velilla and E. Martinez-Salas, 2015 Gemin5: A Multitasking RNA-Binding Protein Involved in Translation Control. Biomolecules 5: 528-544.

Praveen, K., Y. Wen, K. M. Gray, J. J. Noto, A. R. Patlolla et al., 2014 SMA-causing missense mutations in survival motor neuron (Smn) display a wide range of phenotypes when modeled in Drosophila. PLoS Genet 10: e1004489.

Praveen, K., Y. Wen and A. G. Matera, 2012 A Drosophila model of spinal muscular atrophy uncouples snRNP biogenesis functions of survival motor neuron from locomotion and viability defects. Cell Rep 1: 624-631.

Raimer, A. C., K. M. Gray and A. G. Matera, 2017 SMN - A chaperone for nuclear RNP social occasions? RNA Biol 14: 701-711.

Ryder, E., M. Ashburner, R. Bautista-Llacer, J. Drummond, J. Webster et al., 2007 The DrosDel deletion collection: a Drosophila genomewide chromosomal deficiency resource. Genetics 177: 615-629.

Sauterer, R. A., R. J. Feeney and G. W. Zieve, 1988 Cytoplasmic assembly of snRNP particles from stored proteins and newly transcribed snRNA's in L929 mouse fibroblasts. Exp Cell Res 176: 344-359.

Schrank, B., R. Gotz, J. M. Gunnersen, J. M. Ure, K. V. Toyka et al., 1997 Inactivation of the survival motor neuron gene, a candidate gene for human spinal muscular atrophy, leads to massive cell death in early mouse embryos. Proc Natl Acad Sci U S A 94: 9920-9925.

Sen, A., D. N. Dimlich, K. G. Guruharsha, M. W. Kankel, K. Hori et al., 2013 Genetic circuitry of Survival motor neuron, the gene underlying spinal muscular atrophy. Proc Natl Acad Sci U S A 110: E2371-2380.

Shababi, M., C. L. Lorson and S. S. Rudnik-Schoneborn, 2014 Spinal muscular atrophy: a motor neuron disorder or a multi-organ disease? J Anat 224: 15-28.

Shin, E. M., H. S. Hay, M. H. Lee, J. N. Goh, T. Z. Tan et al., 2014 DEAD-box helicase DP103 defines metastatic potential of human breast cancers. J Clin Invest 124: 3807-3824.

Shpargel, K. B., and A. G. Matera, 2005 Gemin proteins are required for efficient assembly of Sm-class ribonucleoproteins. Proc Natl Acad Sci U S A 102: 17372-17377.

Starheim, K. K., D. Gromyko, R. Evjenth, A. Ryningen, J. E. Varhaug et al., 2009 Knockdown of human N alpha-terminal acetyltransferase complex C leads to p53-dependent apoptosis and aberrant human Arl8b localization. Mol Cell Biol 29: 3569-3581.

Sumner, C. J., and T. O. Crawford, 2018 Two breakthrough gene-targeted treatments for spinal muscular atrophy: challenges remain. J Clin Invest 128: 3219-3227.

Tizzano, E. F., and R. S. Finkel, 2017 Spinal muscular atrophy: A changing phenotype beyond the clinical trials. Neuromuscul Disord 27: 883-889.

Van Damme, P., M. Lasa, B. Polevoda, C. Gazquez, A. Elosegui-Artola et al., 2012 N-terminal acetylome analyses and functional insights of the N-terminal acetyltransferase NatB. Proc Natl Acad Sci U S A 109: 12449-12454.

Varland, S., C. Osberg and T. Arnesen, 2015 N-terminal modifications of cellular proteins: The enzymes involved, their substrate specificities and biological effects. Proteomics 15: 2385-2401.

Wang, J., and G. Dreyfuss, 2001 A cell system with targeted disruption of the SMN gene: functional conservation of the SMN protein and dependence of Gemin2 on SMN. J Biol Chem 276: 9599-9605.

West, S., N. J. Proudfoot and M. J. Dye, 2008 Molecular dissection of mammalian RNA polymerase II transcriptional termination. Mol Cell 29: 600-610.

Xu, C., H. Ishikawa, K. Izumikawa, L. Li, H. He et al., 2016 Structural insights into Gemin5-guided selection of pre-snRNAs for snRNP assembly. Genes Dev 30: 2376-2390.

Yan, X., J. F. Mouillet, Q. Ou and Y. Sadovsky, 2003 A novel domain within the DEAD-box protein DP103 is essential for transcriptional repression and helicase activity. Mol Cell Biol 23: 414-423.

Yang, J., P. J. Fuller, J. Morgan, H. Shibata, C. D. Clyne et al., 2015 GEMIN4 functions as a coregulator of the mineralocorticoid receptor. J Mol Endocrinol 54: 149-160.

Yong, J., M. Kasim, J. L. Bachorik, L. Wan and G. Dreyfuss, 2010 Gemin5 delivers snRNA precursors to the SMN complex for snRNP biogenesis. Mol Cell 38: 551-562.

Youkharibache, P., S. Veretnik, Q. Li, K. A. Stanek, C. Mura et al., 2018 The Small β-barrel Domain: A Survey-based Structural Analysis. bioRxiv: doi: https://doi.org/10.1101/140376

